# Zika Virus Infection Prevents Host mRNA Nuclear Export by Disrupting UPF1 Function

**DOI:** 10.1101/2020.12.03.410837

**Authors:** Kristoffer Leon, Ryan Flynn, Mir M. Khalid, Krystal A. Fontaine, Tom Nguyen, G. Renuka Kumar, Camille R Simoneau, Sakshi Tomar, David Jimenez-Morales, Mariah Dunlap, Julia Kaye, Priya S Shah, Steven Finkbeiner, Nevan J Krogan, Carolyn Bertozzi, Jan E. Carette, Melanie Ott

**Author notes:** Address correspondence to, Gladstone Institutes, 1650 Owens Street, San Francisco, CA 9415.

## Abstract

Zika virus (ZIKV) is a mosquito-borne RNA virus that can infect fetuses *in utero* causing characteristic neurodevelopmental disorders including microcephaly. We previously showed that ZIKV infection downregulates expression of up-frameshift protein 1 (UPF1), a helicase/ATPase and central regulator of the nonsense-mediated mRNA decay pathway. Here, we identify a novel function of nuclear UPF1 in mRNA export. Using crosslinking immunoprecipitation of UPF1 followed by sequencing of associated transcripts as well as fluorescence in situ hybridization experiments, we find widespread mRNA accumulation in the nucleus of human neural progenitor cells (NPCs) upon ZIKV infection or UPF1 knockdown. Knockdown of *FREM2*, a top UPF1 target transcript encoding an extra-cellular matrix protein critical in fetal development, decreased expression of pluripotency markers and increased expressed neuronal differentiation in NPCs, consistent with the model that trapping *FREM2* mRNA in the nucleus perturbs proper NPC function. Collectively, our data uncover a new posttranscriptional mechanism by which ZIKV “shuts off” host mRNA export via UPF1. As we find UPF1 linked to many neurodevelopment pathways, we propose that the lack of host mRNA export contributes to the neurodevelopmental defects associated with ZIKV infection.

## Introduction

Zika virus (ZIKV) is a mosquito-borne, enveloped virus with a positive sense, single stranded RNA genome (Musso & Gubler, 2016). ZIKV is a member of the *Flaviviridae* family that includes Dengue virus (DENV), West Nile virus (WNV) and Hepatitis C virus (HCV). In 2015, an outbreak of ZIKV in Brazil was linked to a dramatic increase in the number of infants born with microcephaly (Fauci & Morens, 2016; Pierson & Diamond, 2018). It was subsequently shown that *in utero* ZIKV infects neural progenitor cells (NPCs), resulting in neurodevelopmental delays that ultimately cause a range of birth defects including microcephaly, ocular damage, and contractures, collectively known as congenital Zika syndrome (Costa et al., 2016; Cugola et al., 2016; Shao et al., 2016; Souza et al., 2016; Tang et al., 2016). NPCs are critical for brain development as they differentiate into the glial and neuronal cells that compose the majority of the brain parenchyma (Martínez-Cerdeño & Noctor, 2018). Proposed molecular mechanisms by which ZIKV disrupts NPC function and differentiation include perturbation of ANKLE2/VRK1, centrosomal organization, autophagy, apoptosis and unfolded protein response pathways (Link et al., 2019; Ojha et al., 2018; Saade et al., 2020; Shah et al., 2018; Wen et al., 2019). However, it remains unknown how ZIKV manipulates multiple cellular pathways at once to cause such widespread developmental reprogramming.

We and others previously showed that ZIKV infection suppressed the host nonsense-mediated mRNA decay (NMD) pathway (Fontaine et al., 2018; M. Li et al., 2019). NMD is an RNA quality control mechanism that targets faulty host transcripts for degradation and acts as an antiviral pathway on many viral species, particularly single-stranded RNA viruses (Balistreri, Bognanni, & Mühlemann, 2017; Leon & Ott, 2020). Different models of the NMD pathway have been described, but the exon-junction complex (EJC)-mediated NMD pathway is the best defined (Kurosaki & Maquat, 2016; Kurosaki, Popp, & Maquat, 2019). This pathway is classically triggered by premature termination codons in mRNAs, but can be also be initiated by normal 3’UTRs (Niels H. Gehring, Neu-Yilik, Schell, Hentze, & Kulozik, 2003).

EJCs are normally loaded onto mRNAs in the nucleus (Popp & Maquat, 2013); the RNAs are then exported to the cytoplasm to undergo translation, leading to proper displacement of the EJCs (N. H. Gehring, Lamprinaki, Kulozik, & Hentze, 2009). Non-displaced EJCs recruit the master regulator and helicase up-frameshift protein 1 (UPF1), which induces a recruitment cascade activating NMD-mediated degradation of the faulty transcript (Loh, Jonas, & Izaurralde, 2013; Lykke-Andersen et al., 2014). However, the NMD pathway and UPF1 also target non-faulty mRNAs, especially those with long and GC-rich 3’ untranslated regions (UTRs), a feature found in many viral RNAs (Imamachi, Salam, Suzuki, & Akimitsu, 2017; Kebaara & Atkin, 2009; Peccarelli & Kebaara, 2014; Toma, Rebbapragada, Durand, & Lykke-Andersen, 2015). Because knockdown of UPF1 enhances replication of many RNA viruses including ZIKV, WNV, DENV, Rous Sarcoma virus, Potato virus X, Pea Enation Mosaic virus 2, and Turnip Crinkle virus, NMD is recognized as a *bone fide* antiviral restriction pathway, often targeted by viruses including ZIKV for inactivation (Leon & Ott, 2020).

ZIKV inactivation of the antiviral activity of NMD through interactions between the ZIKV capsid protein and UPF1 has been previously described (Fontaine et al., 2018; M. Li et al., 2019). Similar interactions of capsid proteins with the NMD pathway were subsequently confirmed in other flaviviruses (M. Li et al., 2019). However, ZIKV is unique in that infection or ZIKV capsid expression selectively downregulated UPF1 expression in the host nucleus (Fontaine et al., 2018), which is unusual as viral RNA replication occurs exclusively in the host cytoplasm. Thus, it remains unclear why nuclear UPF1 is targeted during ZIKV infection.

Here we show that UPF1 occupancy of host transcripts is globally decreased during infection with many of these transcripts downregulated in the cytoplasm. ZIKV infection or UPF1 knockdown resulted in polyadenylated transcript accumulation in the nucleus, causing decreased protein levels of FREM2, a UPF1 target, and consequent perturbation of NPC differentiation. We propose that by targeting nuclear UPF1 and trapping host mRNAs in the nucleus, ZIKV has evolved a mechanism to “shut off” host mRNA function while promoting translation of its own proteins. This mechanism describes a new central role for UPF1 in mRNA export connected to many cellular pathways associated with neurodevelopment.

## Results

### ZIKV infection decreases UPF1 interaction with the 3’UTR of host transcripts

While ZIKV inhibits NMD by capsid-mediated degradation of UPF1, the functional impact of UPF1 loss is still unknown. We first aimed to define the UPF1-RNA binding landscape by performing infrared crosslinking immunoprecipitation and RNA sequencing (irCLIP-Seq) in human induced pluripotent stem cell-derived NPCs (Zarnegar et al., 2016). UPF1 is an RNA helicase and ATPase found associated with many transcripts (Fiorini, Bagchi, Le Hir, & Croquette, 2015). To identify differences in UPF1 binding during infection, NPCs were mock-infected or infected with ZIKV (isolate PRVABC59) at an MOI of 1 for 48 hours, followed by UV-crosslinking of transcripts with proteins and immunoprecipitation of UPF1 (**Figure 1A**). RNA-protein complexes were separated by SDS-PAGE, and mass spectrometry was performed on the excised bands to confirm UPF1 enrichment (**Figure S1A, Supplemental File 1**) before RNA was extracted and submitted to next-generation sequencing (**Figure 1A, Supplemental File 2**).

**Figure 1:**
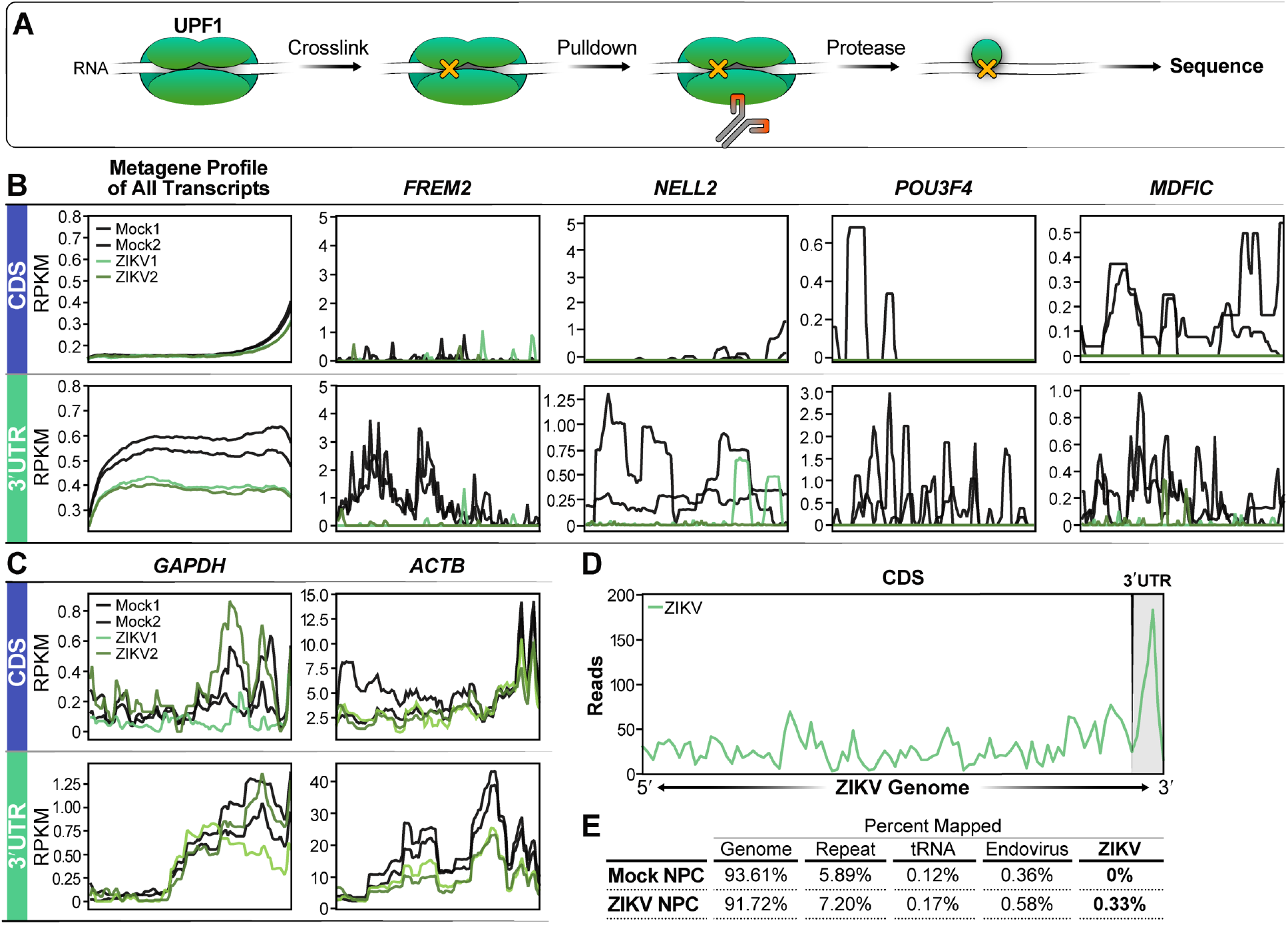
UPF1-host transcript interactions are decreased during ZIKV infection of NPCs. **A)** irCLIP schematic describing the workflow to obtain sequencing data. After 48 hrs of ZIKV infection, UPF1 and RNA are crosslinked using UV light, followed by UPF1 pulldown and protease degradation to expose the UPF1-bound RNA for sequencing. **B)** Metagene profile of all transcripts sequenced from the irCLIP experiment. The graphs show Reads Per Kilobase of transcript, per Million mapped reads (RPKM) values for positions in the CDS and the 3’UTR. Experiment was produced from 2 biological replicates. Representative metagene plots (FREM2, NELL2, POU3F4, MDFIC) of loci found to have a loss of UPF1 interaction. **C)** A metagene plot of GAPDH, a transcript resistant to UPF1 degradation, and ACTB are shown as negative controls. **D)** Metagene plot of reads mapping to the ZIKV genome, with the 3’UTR marked. **E)** Tabular breakdown of read map percentages from the UPF1-CLIP experiment.

In mock-infected NPCs, UPF1 bound 6778 transcripts, predominantly at the 3’UTR, as previously reported (Kebaara & Atkin, 2009; Toma et al., 2015). In ZIKV-infected NPCs, this number decreased to 4557 transcripts, with a marked decrease in binding at the 3’UTR (**Figure 1B**). Overall, a greater than 4-fold decrease in binding was observed in 3% of UPF1 target transcripts. Notably, known NMD-resistant transcripts such as glyceraldehyde-3-phosphate dehydrogenase (GAPDH) maintained UPF1 interaction, highlighting that ZIKV infection did not impact all transcripts uniformly (**Figure 1C**).

In infected samples, UPF1 also interacted with ZIKV RNA, again mostly within the 3’UTR, supporting the model that UPF1 targets the viral RNA to restrict infection (**Figure 1D**). To determine if viral RNA sequestered UPF1 away from host RNA, leading to decreased UPF1 occupancy during infection, we quantified the percent of UPF1-bound reads mapping to host vs. viral RNAs. The majority of sequences isolated with UPF1 pulldown mapped to the host transcriptome; viral reads only accounted for 0.33% of all mapped reads, excluding a “sponge” effect of viral RNA for UPF1 (**Figure 1E**).

### Selective reduction of UPF1 target transcripts in the cytoplasm of infected NPCs

During ZIKV infection, we saw a widespread decrease in UPF1 occupancy of host transcripts. To understand the consequences of the loss of UPF1, we sought to define the changes in the abundance of transcripts identified in the irCLIP-Seq experiment. We performed whole transcriptome sequencing of ZIKV-infected NPCs followed by differential expression analysis (**Supplemental File 3**). As expected, genes associated with the interferon response were predominantly upregulated upon infection, consistent with previous RNA-seq studies of ZIKV-infected NPCs (C. Li et al., 2016; Liu et al., 2019) (**Figure 2A, Supplemental Figure 2C**). The NMD pathway and UPF1 are associated with RNA degradation, and we previously found that ZIKV infection upregulated select NMD target mRNAs (Fontaine et al., 2018). We thus expected that transcripts that had lost UPF1 interaction upon infection would be selectively upregulated. However, the abundance of transcripts which lost UPF1 occupancy upon infection was not significantly changed, indicating that the majority of UPF1 targets were not automatic subjects of NMD (**Figure 2B, Supplemental File 2**). Metascape analysis of transcripts with stable abundance but loss of UPF1 binding showed a characteristic enrichment for neural and neurodevelopmental functions, underscoring the relevance of UPF1 binding for proper neurodevelopment. Axon guidance, which is essential for appropriate establishment of neural circuits in the brain, represented the top pathway identified (**Figure 2C**) (Stoeckli, 2018).

**Figure 2:**
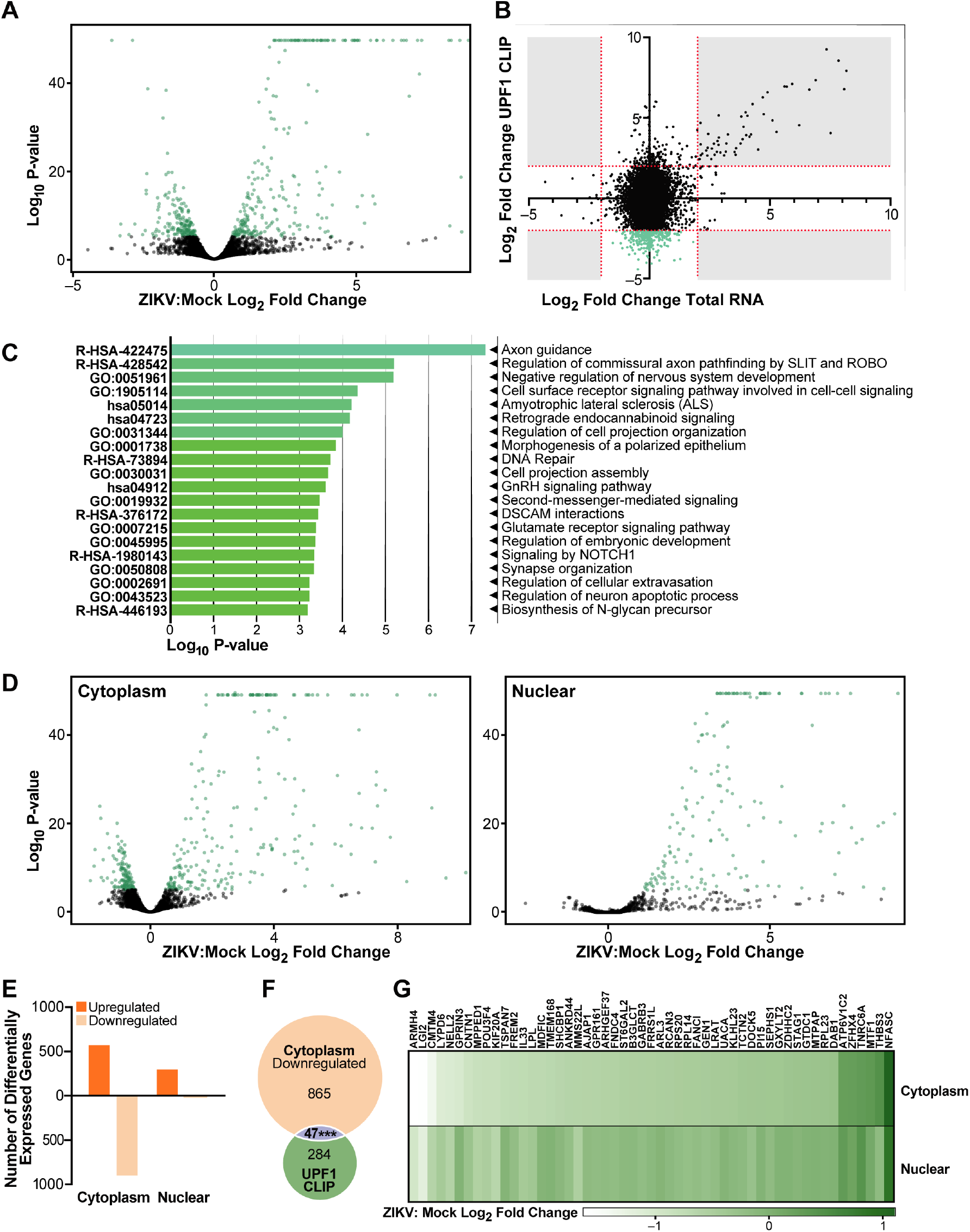
Loss of UPF1 interaction during ZIKV infection causes mRNA downregulation in the cytoplasm. **A)** Volcano plot of whole transcriptome RNA sequencing of ZIKV infected NPCs compared to Mock, n=2 biological replicates. **B)** X-axis shows ZIKV: Mock log_2_ fold change in whole transcriptome sequencing, Y-Axis shows ZIKV:Mock log_2_ fold change from the UPF1 CLIP experiment. The green transcripts are >2 log_2_ fold change in the UPF1-CLIP and <2 log_2_ fold change in total RNA. **C)** Metascape analysis of transcripts identified to have a significant loss of UPF1 interaction in ZIKV infected cells compared to Mock. **D)** Volcano plot of sequencing from ZIKV-infected NPCs compared to Mock fractionated into cytoplasmic and nuclear fractions, n=2 biological replicates. **E)** Number of differentially expressed genes from sequencing of ZIKV-infected NPCs compared to Mock fractionated into cytoplasmic and nuclear fractions. **F)** Overlap of transcripts found to be significantly downregulated in the cytoplasm and those with greater than a 2 log_2_ fold decrease in the UPF1 CLIP. Statistics performed by hypergeometric probability test, ***, P ≤ 0.001. **G)** A heatmap showing log_2_ fold changes of transcripts identified in the CLIP experiment in cytoplasmic and nuclear fractions.

Since transcript abundance was not affected by UPF1 occupancy, we next hypothesized the loss of UPF1 binding may instead impact the subcellular localization of cellular mRNAs. This was based on the knowledge that UPF1 translocates between the nucleus and cytoplasm (Varsally & Brogna, 2012), and our previous observation that UPF1 is selectively downregulated in the cell nucleus upon infection (Fontaine et al., 2018). To test this hypothesis, we fractionated ZIKV-infected NPCs into cytoplasmic and nuclear compartments and performed RNA sequencing and differential expression analysis with each fraction (**Supplemental File 4,5**). We confirmed successful fractionation by comparing the log_2_ fold change of known cytoplasmic (*GAPDH*) and nuclear (*ANRIL*) transcripts, which showed 3.2-fold enrichment of *GAPDH* in the cytoplasmic fraction and a 2.3-fold increase of *ANRIL* in the nuclear fraction, as expected (**Supplemental Figure 2A**). Upon ZIKV infection, we identified 585 and 312 significantly upregulated mRNAs in the cytoplasm and nucleus, respectively (**Figure 2D**). These upregulated transcripts were predominantly interferon response genes (**Supplemental Figure 2D**). In contrast, downregulated genes were overwhelmingly cytoplasmic, with 912 genes downregulated in the cytoplasm compared to 22 in the nuclear fraction (**Figure 2E**). A significant overlap was found between the 912 transcripts with reduced cytoplasmic expression and the 331 transcripts that lost UPF1 occupancy without changing overall abundance (**Figure 2F**). In contrast, considerably less overlap existed with transcripts that were upregulated in the cytoplasm and had lost UPF1 binding (**Supplemental Figure 2B**). Moreover, of the top 53 genes with the most significant decrease in UPF1 interaction upon infection, all but 6 were exclusively downregulated in the cytoplasm (**Figure 2G**). These results show a selective loss of UPF1 target mRNAs in the cytoplasm of infected cells. As total abundance of these transcripts was unchanged, we considered the possibility that they accumulate in the nucleus upon ZIKV infection.

### ZIKV infection, ZIKV capsid expression or UPF1 knockdown results in accumulation of host transcripts in the nucleus

To test the hypothesis that transcripts were selectively accumulating in the nucleus in ZIKV-infected cells, we analyzed global transcript localization using RNAscope analysis with a polyA tail probe. We first optimized the technology in Huh7-Lunet cells, a hepatoma cell line frequently used to study flavivirus infection (Vicenti et al., 2018), and NPCs, both infected with ZIKV at an MOI of 1. In both cell types, polyadenylated RNA fluorescence was significantly increased in the nucleus upon infection, supporting the model that mRNA location rather than abundance is perturbed in ZIKV-infected cells (**Figure 3A,B**). To understand if this effect was UPF1-mediated, we leveraged the previously demonstrated ability of ZIKV capsid to specifically degrade nuclear UPF1 (Fontaine et al., 2018), and used Huh7-Lunet cells expressing ZIKV capsid protein in a tetracycline-inducible manner. Induction of capsid expression by addition of doxycycline led to reduced levels of nuclear UPF1 (**Figure 3C**). This also led to a similar accumulation of polyadenylated RNAs in the cell nucleus, mirroring the effect observed in ZIKV-infected cells (62% nuclear fluorescence in vector vs 78% in capsid overexpressing cells) (**Figure 3D**). Furthermore, siRNA-mediated knockdown of UPF1 in NPCs also resulted in retention of polyadenylated RNAs in the nucleus (36% nuclear fluorescence in control vs 40% in siUPF1 NPCs) (**Figure 3E, Supplemental Figure 3**). Collectively, these data indicate that UPF1 regulates nuclear mRNA export, a function perturbed upon nuclear UPF1 degradation during ZIKV infection.

**Figure 3:**
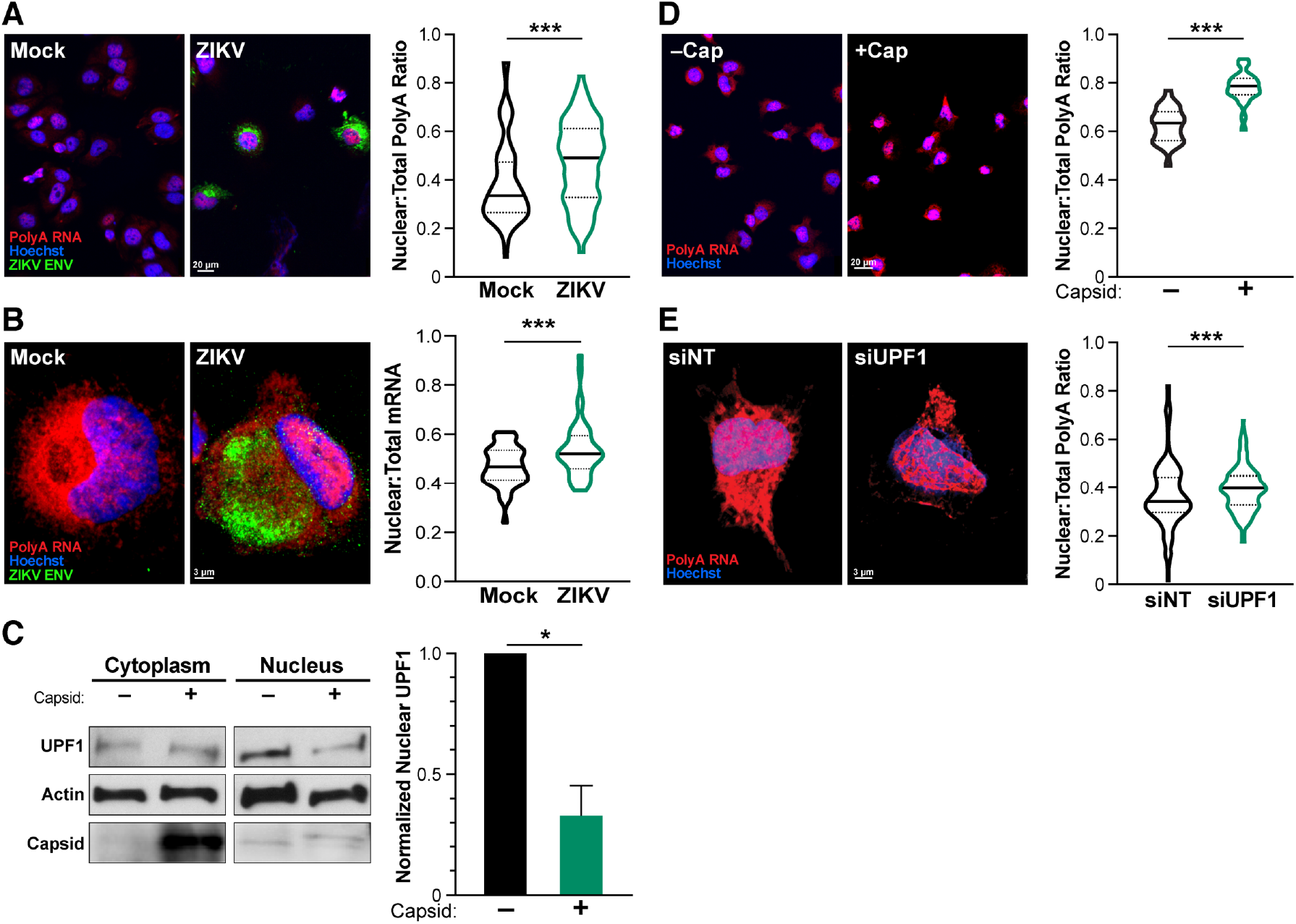
ZIKV-mediated degradation of UPF1 leads to mRNA retention in the nucleus. **A and B)** During ZIKV infection (ZIKV ENV, green), polyA RNA (red) is shifted toward the nucleus (Hoechst, blue) in Huh7-Lunet cells (A) and NPCs (B). Statistics produced by a Linear Mixed Model. 3 biological replicates, n=25 cells (A) or 15 cells (B) per condition per replicate. **C)** Tetracycline-inducible Capsid expression in Huh7-Lunet cells was used to degrade nuclear UPF1. Leptomycin B (LMB) was used to increase UPF1 degradation. Statistics performed by Student’s t-test, n=3, and representative western blot shown. **D)** Capsid overexpression in Huh7-Lunet cells results in an increased ratio of polyA RNAs (red) in the nucleus. N=3 biological replicates, 25 cells per condition per replicate. **E)** Knockdown of UPF1 in NPCs results in an increased ratio of polyA RNAs (red) in the nucleus. NPCs were treated with siNT and siUPF1 for 96 hours. Statistics produced by a Linear Mixed Model. n=3 biological replicates, 15 cells per condition per replicate. *, P ≤ 0.05; **, P ≤ 0.01; ***, P ≤ 0.001. Error bars are SEM.

### Depletion of UPF1 leads to nuclear retention of *FREM2* mRNA, depletion of FREM2 protein, and alterations to NPC differentiation markers

Next, we focused on *FREM2*, the transcript with the largest fold decrease in UPF1 interaction in the irCLIP studies (**Figure 1B**). FREM2 is an extracellular matrix protein involved in cell-cell interactions that is important in many developmental pathways including tissue and vascular morphogenesis (Pavlakis, Chiotaki, & Chalepakis, 2011; Timmer, Mak, Manova, Anderson, & Niswander, 2005). The *FREM2* transcript was also specifically downregulated in the cytoplasm of infected NPCs (**Figure 2G**). We confirmed selective cytoplasmic downregulation of *FREM2* mRNA expression upon infection by qPCR (**Figure 4A**). Using RNAscope, we found enrichment of the *FREM2* transcript in the nucleus of cells treated with UPF1-targeting siRNAs as compared to cells treated with non-targeting control siRNAs, where it was found in both the cytoplasm and the nucleus (**Figure 4B**). FREM2 protein levels were significantly decreased in UPF1 siRNA-treated cells (42% decrease in siUPF1 compared to the siNT control), supporting the model that nuclear mRNA retention leads to reduced translation of UPF1 target transcripts during ZIKV infection (**Figure 4C**).

**Figure 4:**
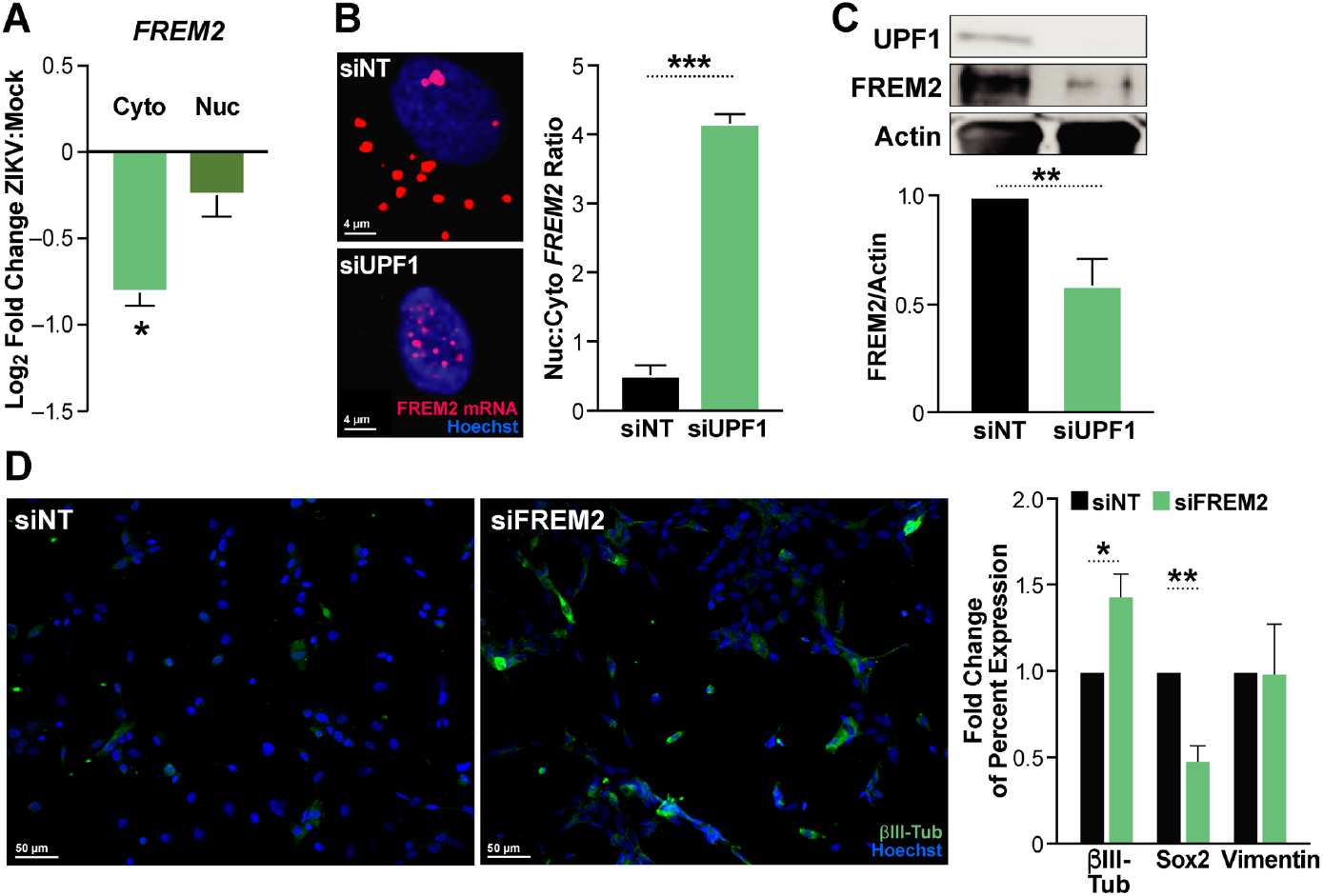
UPF1 knockdown leads to retention of *FREM2* mRNA and decreased protein production, which alters differentiation markers. **A)** qPCR of *FREM2* in cytoplasmic and nuclear fractions in ZIKV infected NPCs. **B)** RNAscope of *FREM2* in UPF1 knockdown NPCs. NPCs were treated with siRNAs for 96 hours prior to harvesting for microscopy. Number of nuclear puncta compared to cytoplasmic as calculated by Imaris. 3 biological replicates, 10 cells per biological replicate per condition averaged. Statistics produced by Student’s t-test. **C)** Western blot of siNT and siUPF1 treated NPCs. NPCs were treated with siRNAs for 96 hours prior to harvesting for microscopy. Densitometric analyses of FREM2 were performed using ImageJ to quantify relative band intensities. **D)** siNT and siFREM2 treated NPCs stained for βIII-Tubulin, Soxs2 and Vimentin. NPCs were treated with siRNAs for 7 days prior to analysis. Statistics produced by Student’s t-test. *, P ≤0.05; **, P ≤0.01; ***, P ≤ 0.001. Error bars are SEM.

To determine the functional consequences of reduced FREM2 expression, we used siRNAs to knockdown FREM2 in NPCs (**Supplemental Figure 4**). FREM2 knockdown increased the percentage of cells expressing the neuronal marker βIII-Tubulin by 44%, and decreased the percentage of cells expressing the pluripotency marker Sox2 by 51% (**Figure 4D**). These results indicate that by preventing FREM2 nuclear export and protein expression, ZIKV infection may promote premature neuronal differentiation of NPCs, thus reducing the number of NPCs that determine brain and ultimately head size.

## Discussion

In this study, we identified a novel mechanism by which ZIKV infection prevents RNA export from the nucleus, a phenotype described in other viral infections, but not seen previously with flaviviruses (Fortes, Beloso, & Ortín, 1994; Kumar & Glaunsinger, 2010; Kuss, Mata, Zhang, & Fontoura, 2013; Sakuma et al., 2014). Many transcripts lost UPF1 occupancy during ZIKV infection, especially in the 3’UTR, consistent with the finding that UPF1 expression is downregulated in ZIKV infection. Loss of UPF1 binding altered transcript localization with only a minor effect on transcript abundance, underscoring the functional significance of the role of UPF1 in mRNA export. The accumulation of mRNA in the nucleus induced by ZIKV infection could be recapitulated both by ZIKV capsid expression, which is known to degrade nuclear UPF1, and by UPF1 knockdown, linking the new function of UPF1 with its nuclear localization. Lastly, we show that when UPF1 is unable to export *FREM2* mRNA from the nucleus, FREM2 protein abundance decreases, which may disrupt neural differentiation. The majority of these experiments were performed in human NPCs, the natural target cells of ZIKV in fetuses, providing a physiologically relevant view into pathways and transcripts disrupted by ZIKV infection and how that might lead to congenital Zika syndrome.

UPF1 is an RNA helicase involved with a plethora of functions within the cell. Besides its function in NMD, UPF1 also plays important roles in several other RNA decay pathways (Kim & Maquat, 2019), as well as telomere maintenance (Chawla et al., 2011), and ubiquitination (Feng, Jagannathan, & Bradley, 2017). While this study was underway, a study in fruit flies connected nuclear UPF1 function with co-transcriptional binding and export of mRNAs from the nucleus (Singh et al., 2019). UPF1 is also required for HIV RNA export from the nucleus (Ajamian et al., 2015).

Here we show a role of nuclear UPF1 in mRNA export in mammalian cells, which is disrupted by ZIKV infection. While the precise mechanisms of UPF1-mediated nuclear export are not yet clear, two findings underscore its significance: 1) most transcripts were not upregulated in NPCs after loss of UPF1 binding, which points to another major function of UPF1 in cells other than NMD, and 2) many polyadenylated transcripts were retained in the nucleus upon UPF1 knockdown, demonstrating that the effect of UPF1 on mRNA export is widespread. Additional studies are needed to determine whether specific features of these mRNAs, such as the 3’UTR sequence or structure, underlie their targeting for export by UPF1 (Carmody & Wente, 2009). Similarly, additional studies are needed to determine the features that lead to UPF1 enrichment in the ZIKV 3’UTR. Interestingly, a non-coding subgenomic flavivirus RNA (SfRNA) is derived from the 3’ UTR of ZIKV and is known to be resistant to XRN1 degradation (Göertz, Abbo, Fros, & Pijlman, 2018). The production of an SfRNA is shared among flaviviruses and antagonizes the interferon response. UPF1 potentially interacts with ZIKV SfRNA, which could play an important role in NMD evasion.

The host shutoff effect is a common viral strategy to reduce host gene expression, potentially for the purposes of competing with cellular resources and evading immune responses. Flaviviruses have been reported to cause host shutoff by translational repression (Roth et al., 2017). The host shutoff described here is achieved by retention of host transcripts in the nucleus. We confirm that a single viral protein, the capsid protein, is sufficient to specifically degrade nuclear UPF1 and cause nuclear mRNA retention. This underscores the importance of the localization of ZIKV capsid to the cell nucleus (Oliveira, Mohana-Borges, de Alencastro, & Horta, 2017), which is independent from its function as a structural component of the virion as the virus replicates exclusively outside of the nucleus (Mohd Ropidi, Khazali, Nor Rashid, & Yusof, 2020). Our study also highlights the significance of nuclear UPF1, which is less studied than cytoplasmic UPF1, although it has been associated with cell cycle progression, DNA replication, telomere maintenance, and mRNA release (Singh et al., 2019; Varsally & Brogna, 2012).

Nuclear retention of host mRNAs caused by the disruption of UPF1 function could contribute to the neurodevelopmental defects seen during fetal ZIKV infections. Since UPF1 depletion impacts many mRNAs, it may represent a central mechanism explaining pleotropic effects of ZIKV infection on diverse cellular pathways. We pursued *FREM2* as a top UPF1 target, and show that it is downregulated at the protein level when UPF1 is unable to perform its role as a nuclear mRNA export regulator. FREM2 is both a member of the FRAS/FREM complex and regulates its formation (Pavlakis et al., 2011). The FRAS/FREM complex is found in cellular basement membranes and is expressed differentially during development (Timmer et al., 2005). FREM2 itself is important for proper development of the eye and FREM2 mutation is associated with Fraser syndrome, in which cryptopthalamos (a congenital defect where the eyes are covered completely by skin and often associated with small or missing eyeballs) is commonly seen (Slavotinek & Tifft, 2002). As developmental defects in the eyes are also seen in children with congenital Zika syndrome, the retention of the *FREM2* transcript in the nucleus could be directly involved in pathogenesis (de Paula Freitas, Ventura, Maia, & Belfort, 2017; Ventura & Ventura, 2018).

Surprisingly, our studies in NPCs and Huh7 hepatoma cells indicate that at least in these cell types the role of UPF1 in nuclear export affects more mRNAs than its known function in NMD. How this function is regulated and whether it involves known UPF1 interacting proteins such as the EJC or the known RNA export pathways such as CRM-1, remains to be determined. The finding that ZIKV has evolved a mechanism to selectively target nuclear UPF1 and the mRNA export function of the protein underscores its central yet so far underappreciated role in biology.

## Materials and Methods

### Cell Culture and Viruses

Human iPSC-derived NPCs were generated and maintained as described previously (Consortium, 2017). The human fibroblast cell line used to generate iPSCs came from the Coriell Institute for Medical Research and Yale Stem Cell Center. The iPSCs used in these studies was the CTRL2493n17 line. CTRL2493n17 was derived from the parental fibroblast line ND31845 that was biopsied from a healthy female at 71 years of age. iPSCs were cultured and maintained in complete mTESR (StemCell Techologies). NPCs were differentiated and maintained using EFH media (Stemline Neural Stem Cell Media [Sigma Aldrich], EGF [R&D Biosystems], rhFGF basic [R&D Biosystems]. Heparin Sulfate [Sigma Aldrich]). NPCs were dissociated and plated onto Matrigel-coated plates (Corning) prior to infecting with ZIKV (MOI of 1) or treating with siRNAs. Experiments were harvested 48 hours post infection and up to 7 days after siRNA treatment. Huh7-Lunet cells (Ralf Bartenschlager, Heidelberg University), and Vero cells were maintained in Dulbecco’s modified Eagle’s medium (DMEM) with 10% fetal bovine serum (FBS), 2 mM l-glutamine, 100 U/ml penicillin, and 100 μg/ml streptomycin.

N-terminally Strep TagII tagged ZIKV Capsid was cloned to pLVX-TetOne-Puro Vector (Clontech, Cat: 631849) using BamHI & EcoRI cut sites. We used a 2nd generation lentiviral system to generate a stably expressing ZIKV Capsid Huh7-Lunet cell line. After transduction, Huh7-Lunet cells were selected with puromycin (2ug/ml) for one weeks and later clonally expanded. Clones with the highest and specific expression were used for the experiment.

The strain PRVABC59 of ZIKV (ATCC VR-1843) was used for all experiments. ZIKV stocks were propagated in Vero cells (ATCC), and titers were determined by plaque assays on Vero cells. ZIKV infections were performed by adding viral inoculum to DMEM with 2% FBS or EFH followed by a two-hour incubation at 37C with a rock every 15 minutes. After infection was completed, inoculum was aspirated and then fresh DMEM with 10% FBS or EFH was added to the cells. Infected cells were cultured for 48 hours prior to harvesting for all sequencing and IF experiments.

### Antibodies and other reagents

Primary antibodies used were anti-UPF1 (Bethyl laboratories, A300-38A, CST, 12040S and Abcam, ab109363), anti-FREM2 (Invitrogen, PA5-20982), anti-FLAG (Abcam, ab18230), anti-Actin (CST, 4967S), anti-GFAP (Abcam, ab53554), anti-Nestin (Abcam, ab22035) and anti-Beta III tubulin (Abcam, ab18207). Secondary antibodies used include goat anti-rabbit Alexa 488 (Invitrogen, A-11008), goat anti-mouse Alexa 488 (Invitrogen, A-11001), goat anti-rabbit Alexa 594 (Invitrogen, A-11012), goat anti-mouse Alexa 594 (Invitrogen, A-11005), donkey anti-goat Alexa 647 (Invitrogen, A-21447), donkey antirabbit Alexa 488 (Invitrogen, A-21206), donkey anti-mouse Alexa 594 (Invitrogen, A-21203). The RNAScope Multiplex Fluorescent V2 Assay (ACD, 323100) was used with RNAscope Probes include polyA RNA (ACD, 318631) and FREM2 (ACD, 482841). Opal570 (Akoya Biosciences, FP1488001KT) was used for visualization of RNAscope probes. Accell siRNA was used for knockdown of FREM2 (Dharmacon, E-021693-00-0010) and UPF1 (Dharmacon, E-011763-00-0010) according to manufacturer’s instructions. Non-targeting siRNAs used were cat. D-001910-10-20 (Dharmacon). Huh7-Lunets were treated with 60 ng/mL of Leptomycin B (Cayman Chemical) for 16 hours.

### Infrared Crosslinking and Immunoprecipitation

irCLIP was performed as in Zarnegar et al. 2016 (BJ et al., 2016). HeLa cells grown as described above and UV crosslinked to a total of 0.35 J/cm^2^. Whole-cell lysates were generated in CLIP lysis buffer (50 mM HEPES, 200 mM NaCl, 1 mM EDTA, 10% glycerol, 0.1% NP-40, 0.2% Triton X-100, 0.5% N-lauroylsarcosine) and briefly sonicated using a probe-tip Branson sonicator to solubilize chromatin. Each experiment was normalized for total protein amount, typically 1 mg, and partially digested with RNase A (ThermoFisher Scientific, EN0531) for 10 minutes at 37°C and quenched on ice. UPF1 (Bethyl laboratories, A300-38A) IP’s were performed using 15 μg of each antibody with 50 μL Protein G Dynabeads (ThermoFisher Scientific), for 8 hours at 4°C on rotation. Samples were washed sequentially in 1 mL for 1 minute each at 25°C: 1× high stringency buffer (15 mM Tris-HCl, pH 7.5, 5 mM EDTA, 2.5 mM EGTA, 1% Triton X-100, 1% sodium deoxycholate, 120 mM NaCl, 25 mM KCl), 1× high salt buffer (15 mM Tris-HCl pH 7.5, 5 mM EDTA, 2.5 mM EGTA, 1% Triton X-100, 1% sodium deoxycholate, 1 M NaCl), 2× NT2 buffer (50 mM Tris-HCl, pH 7.5, 150 mM NaCl, 1 mM MgCl_2_, 0.05% NP-40). After the NT2 wash, RNA-protein complexes were dephosphorylated with T4 PNK (NEB) for 45 minutes in an Eppendorf Thermomixer at 37°C, 15 seconds 1400rpm, 90 seconds of rest in a 30 μL reaction, pH 6.5, containing 10 units of T4 PNK, 0.1 μL SUPERase-IN (ThermoFisher Scientific), and 6 μL of PEG-400 (16.7% final). Dephosphorylated RNA-protein complexes were then rinsed once with NT2 buffer and 3’-end ligated with T4 RNA Ligase 1 (NEB) overnight in an Eppendorf Thermomixer at 16°C, 15 seconds 1400rpm, 90 seconds of rest in a 60 μL reaction containing 10 units T4 RNA Ligase, 1.5 pmol pre-adenylated-IR800-3’biotin DNA-adapter, 0.1 μL SUPERase-IN, and 6 μL of PEG400 (16.7% final). The following day, samples were again rinsed once with 500 μL NT2 buffer and resuspended in 30μL of 20 mM DTT, 1x LDS (ThermoFisher Scientific) in NT2 buffer. Samples were heated to 75°C for 10 min, and released RNA-protein complexes were separated on 4-12% Bis-Tris SDS-PAGE (1.0mm X 12 well) at 200V for 45 min. Resolved RNP complexes were wet-transferred to nitrocellulose at 550 mA for 45 minutes at 4°C.

Nitrocellulose membranes were imaged using an Odyssey CLx scanner (LiCor), RBP-RNA complexes were excised using scalpels, and RNA was recovered by adding 0.1 mL of Proteinase K reaction buffer (100 mM Tris, pH 7.5, 50 mM NaCl, 1 mM EDTA, 0.2% SDS) and 5 μL of 20mg/mL Proteinase K (ThermoFisher Scientific). Proteins were digested for 60 minutes at 50°C in an Eppendorf Thermomixer. Next, 200 μL of saturated-phenol-chloroform, pH, 6.7 was added to each tube and incubated for 10 minutes at 37°C in an Eppendorf Thermomixer, 1400 rpm. Tubes were briefly centrifuged and the entire contents transferred to a 2 mL Heavy Phase Lock Gel (5Prime, 2302830). Samples were centrifuged for 2 minutes at >13000 rpm. The aqueous layer was re-extracted with 1 mL of chloroform (inverting 10 times to mix; no vortexing) in the same 2 mL Phase Lock Gel tube and centrifuged for 2 minutes at >13000 rpm. The aqueous layer was then transferred to a new 2 mL Heavy Phase Lock Gel tube and extracted again with an additional 1 mL of chloroform. After 2 minutes centrifugation at >13000 rpm, the aqueous layer was transferred to a siliconized 1.5 mL tube and precipitated overnight at −20°C by addition of 10 μL 5M NaCl, 3 μL Linear Polyacrylamide (ThermoFisher Scientific) and 0.8 mL 100% ethanol. RNA fragments were pelleted at >13000 rpm for 45 minutes at 4°C, washed once with 1 mL of ice cold 75% ethanol and air dried.

RNA pellets were resuspended in 12 μL water 1 μL of 3 μM cDNA and 1 μL of 10mM dNTPs and heated to 70°C for 5 minutes then rapidly cooled to 4°C. cDNA Master Mix (4 μL 5x Super Script IV (SSIV) Buffer, 1 μL 100mM DTT, 1 μL SSIV, 6 μL total) was added to the annealed RNA and incubated for 30minutes at 55°C. cDNA:RNA hybrids were captured by addition of 5 μL of MyOne Streptavidin C1 Dynabeads (ThermoFisher Scientific) that had been rinsed and suspended in 50 μL of Biotin-IP buffer (100mM Tris, pH 7.5, 1M NaCl, 1mM EDTA, 0.1% Tween), and end over end rotation for 45 minutes at room temperature. Beads were placed on a 96-well magnet and washed sequentially twice with 100 μL of Biotin IP buffer and 100 μL ice-cold 1xPBS. Beads were resuspended in 10 μL of cDNA elution buffer (8.25 μL water, 1 μL of 1 μM P3 short oligo, and 0.75 μl of 50 mM MnCl_2_) and heated to 95°C for 10 minutes, ramp 0.1 degree/second to 60°C forever. Next 5 μL of circularization reaction buffer was added (3.3 μL water, 1.5 μL 10x Circligase-II buffer, and 0.5 μL of Circligase-II (Epicentre)). cDNA was circularized for 2 hours at 60°C. cDNA was purified with 30 μL of AMPure XP beads (Beckman Coulter) and 75 μL of isopropanol. Samples were incubated for 20 minutes at 25°C, washed twice with 100 μL 80% ethanol, air dried for 5 minutes, and eluted in 14 μL of water. Elution took place at 95°C for 3 minutes and the eluent was immediately transferred to a 96-well magnet. Eluted cDNA was transferred to a new PCR tube containing 15 μL of 2X Phusion HF-PCR Master Mix (NEB), 0.5 μL of 30 μM P3/P6 PCR1 oligo mix and 0.5 μl of 15x SYBR Green I (ThermoFisher Scientific). Real-time quantitative PCR was performed: 98°C 2 min, 15 cycles of 98°C 15 seconds, 65°C 30 seconds, 72°C, 30 seconds, with data acquisition set to the 72°C extension. PCR1 reactions were cleaned up by adding of 4.5 μL of isopropanol, 54 μL of AMPure XP beads and incubation for 10 min. Beads were washed once with 80% ethanol, dried for 5 min, and eluted in 15 μl of water. Illumina flow cell adaptors were added by adding 15 μL 2X Phusion HF-PCR Master Mix and 0.4 μL P3solexa/P6solexa oligo mix and amplified: 98°C 2 min, 3 cycles of 98°C 15 seconds, 65°C 30 seconds, 72°C, 30s seconds. Final libraries were purified by addition of 48 μL of AMPure XP beads and incubation for 5 min. Beads were washed twice with 70% ethanol, dried for 5 min, and eluted in 20 μL of water. 1-2μL of libraries were quantitated by HS-DNA Bioanalyzer. Samples were deep sequenced on the Illumina NextSeq machine: single-end, no index, high-output, 75-bp cycle run. Whole transcriptome RNA sequencing was performed using the methods described above for RNA extraction, library preparation and sequencing.

### Nuclear/cytoplasmic RNA fractionation and sequencing

Fractionation was performed using the Cytoplasmic and Nuclear RNA Purification Kit (Cat. # 2100, Norgen Biotek Corp). Purified RNA was treated with DNase I followed by library preparation using the NuGEN V2 RNA-Sequencing Library Preparation kit (Tecan Genomics). Both RNA and library quality were analyzed via a Bioanalyzer (Agilent). Sequencing was performed on a NextSeq 500 (Illumina): single-end, no index, high-output, 75-bp cycle run.

### IF and RNAscope protocol

For immunofluorescence and RNAscope, infected NPCs were collected at 48 hr and plated onto 22-by 22-mm no. 1.5 coverslips. Cells were then fixed with 4% PFA in PBS for 15 minutes. For the RNAscope protocol, we followed manufacturer’s instructions for adherent cell lines. Briefly, we first dehydrated the cells using 50%, 70% and then 100% ethanol in PBS. This was followed by a rehydration of the cells using 70% and then 50% of ethanol in PBS. Lastly, cells were fully rehydrated in PBS. Cells were then permeabilized by hydrogen peroxide, followed by protease 3 treatment. Next, we hybridized the RNAscope probes to the cells for 2 hours or O/N at 40C. Probe amplification was then performed, followed by labelling with Opal570 (Akoya Biosciences). Nuclei were stained using Hoechst 33258 (Thermofisher).

For immunofluorescence, cells were fixed, permeabilized by 0.1% Triton X-100 (Sigma Aldrich), blocked with 3% bovine serum albumin (Sigma Aldrich) in PBS. Cells were then immunostained with the indicated antibody, followed by the appropriate secondary. Lastly, nuclei were stained with Hoechst 33258.

Microscopy was performed on an LSM880 with Airyscan (Zeiss) or an Olympus FV3000RS. On the LSM880, imaging of Huh7-Lunet cells was performed with 20x magnification objective, while NPCs were imaged with a 63x oil objective. On the FV3000RS, the 20x objective was used for imaging NPCs. All images were taken as a Z-stack.

### Western blot analysis

Cells were lysed in RIPA lysis buffer (50 mM Tris-HCl [pH 8], 150 mM NaCl, 1% NP-40, 0.5% sodium deoxycholate, 0.1% SDS, supplemented with Halt protease inhibitor cocktail [Thermo Fisher Scientific]) to obtain whole-cell lysates or lysed using the NE-PER nuclear and cytoplasmic extraction kit (Thermo Fisher Scientific) to obtain cytoplasmic and nuclear fractions. Proteins were separated by SDS-PAGE and transferred to nitrocellulose membranes (Bio-Rad). Proteins were visualized by chemiluminescent detection with ECL and visualized on a ChemiDoc MP Imaging System (Bio-Rad).

### Computational and Statistical Analyses

For western blot analysis, differences in band intensity were quantified by densitometry using ImageJ (Schneider, Rasband, & Eliceiri, 2012). Student’s t-test was used for statistical analysis of Western Blots. Imaris (Oxford Instruments) was used for analysis of confocal images, using the surface function for polyA mRNA analysis and dot function identifying specific transcripts by RNA-scope. Nuclei were also bounded and identified by the surface function of Imaris. PolyA RNA-scope experiments were statistically analyzed using a linear mixed model to account for individual cell values across multiple biological replicates. Data are represented as means plus standard errors of the means (SEM). Gene set overlap statistics were performed using a hypergeometric test. Statistical significance was defined as follows: *, P ≤0.05; **, P ≤0.01; ***, P ≤ 0.001; ****, P ≤0.0001. Biological replicates are defined as the same experimental design but performed sequentially, with a different cell passage number and on different days.

### Analysis of RNA sequencing data

PCR duplicates were removed using unique molecular identifiers in the RT primer region. The adaptor and barcode sequences were trimmed and reads were mapped step-wise to viral (ZIKV), repetitive and finally non-repetitive (GRCh38) genomes. Bowtie2 indexes were generated using the ‘bowtie2-build’ command in Bowtie2 for the ZIKV (KU501215.1) RNA genome sequences. The specific parameters used for the FAST-iCLIP pipeline were as follows: -f 18 (trims 17 nt from the 5’ end of the read), -l 16 (includes all reads longer than 16 nt), -bm 29 (minimum MAPQ score from bowtie2 of 29 is required for mapping; unique mapping only), -tr 2,3 (repetitive genome) and -tn 2,3 (non-repetitive genome) RT stop intersection (n,m; where n = replicate number and m = number of unique RT stops required per n replicates). Using the -tr/tn 2,3 parameters, a minimum of six RT stops are required to support any single nucleotide identified as a crosslinking site.

Analysis of the sequencing data was performed using a custom analysis pipeline, with the peak finding software uploaded to Github (https://github.com/ChangLab/FAST-iCLIP/tree/lite). Other analyses were performed by aligning the reads to the human genome using STAR, followed by gene counts using Bedtools (AR & IM, 2010). Only reads in the canonical 3’UTR of the human transcripts were counted. The count distribution across the metagene of the CDS and 3’UTRs was visualized using deeptools (F, F, S, BA, & T, 2014). Log_2_ fold changes were calculated by comparing RPKM values between Mock and ZIKV infected cells. Genes with fewer than 10 total read counts were excluded from analysis.

Heat maps and XY plots were produced using Prism 8 (GraphPad).

For total RNA and cytoplasmic/nuclear fractionated sequencing, reads were aligned to the human genome using the STAR aligner (A et al., 2013) (version 2.7.5) followed by HTSeq (S, PT, & W, 2015) (version 0.12.3) to obtain counts and then using the DeSeq2 (MI, W, & S, 2014) (version 1.28.1) pipeline to determine log_2_ fold changes in transcripts. Volcano plots were produced using the R-package, Enhanced Volcano (https://github.com/kevinblighe/EnhancedVolcano).

## Acknowledgements

Stephen Floor, PhD and Michael Wilson, MD provided guidance and mentorship during this project as members of a thesis committee. Library preparation and QCs for sequencing was conducted by Mylinh Bernardi, BS at the Gladstone Genomics Core. The James B. Pendleton Charitable Trust supported the sequencing on the NextSeq 500. Meredith Calvert, PhD and Blaise Ndjamen, PhD of the Gladstone Histology and Light Microscopy core assisted with imaging and image analysis. Reuben Thomas, PhD in the Gladstone Bioinformatics core provided statistical analysis help. Wendy Runyon, BS in the Gladstone Stem Cell core helped with iPSC maintenance. John Carroll provided graphic design expertise, and Kathryn Claiborn, PhD provided manuscript editing.

## Funding

This work was supported by the UCSF Medical Scientist Training Program (KEL), the UCSF Biomedical Sciences Program (KEL), the NINDS/NIH under award F31NS113432 (KEL), the NIAID/NIH under award 5R01AI097552-03 (KEL, MO), NIDA/NIH under award 5DP1DA038043-05 (KEL, MO), NIH/NIAID R01 AI141970 (JEC), NIH/NIAID F32AI112262 (PSS), NIH/NINDS R01 NS101996-01 (SF), NIH/NIAID U19AI1186101 to (NJK), DOD/DARPA HR0011-11-C-0094 (PROPHECY, NJK), the Ottellini Family Discovery Fellowship (KEL), Damon Runyon Cancer Research Foundation DRG-2286-17 (R.A.F.), and the James B. Pendleton Charitable Trust.

## Author Contributions

KEL, MO contributed to experimental design, data acquisition, data analysis and manuscript writing. RAF, CB, JEC contributed to experimental design, data acquisition and data analysis. MK, CS contributed to data acquisition. KAF, TN, ST and GRK contributed to experimental design and data analysis. DJ-M, PSS and NJK contributed with computational analysis. MD, JK and SF contributed assistance with data acquisition.

## Competing Interests

No competing or conflicts of interest to report.

## Supplemental Figures

**Supplemental Figure 1:**
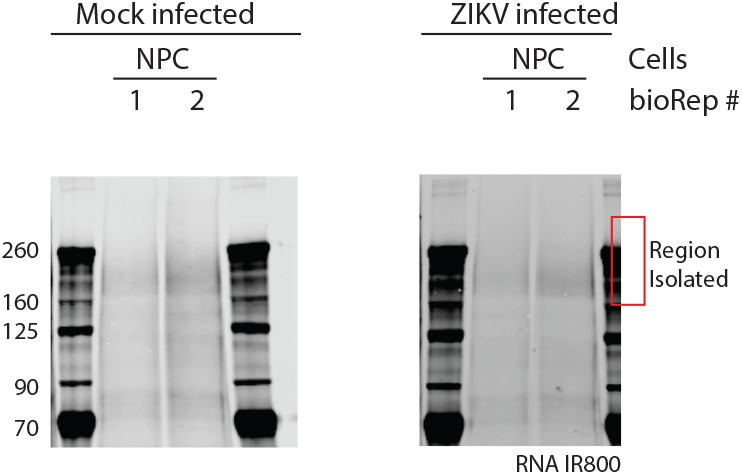
Electrophoresis of RNA pulled down with UPF1, visualized using the IR handle ligated to the RNAs. The region bounded by the red box indicates the part of the gel excised and then analyzed by irCLIP and AP-MS. AP-MS analysis is indicated in Supplemental File 1.

**Supplemental Figure 2.**
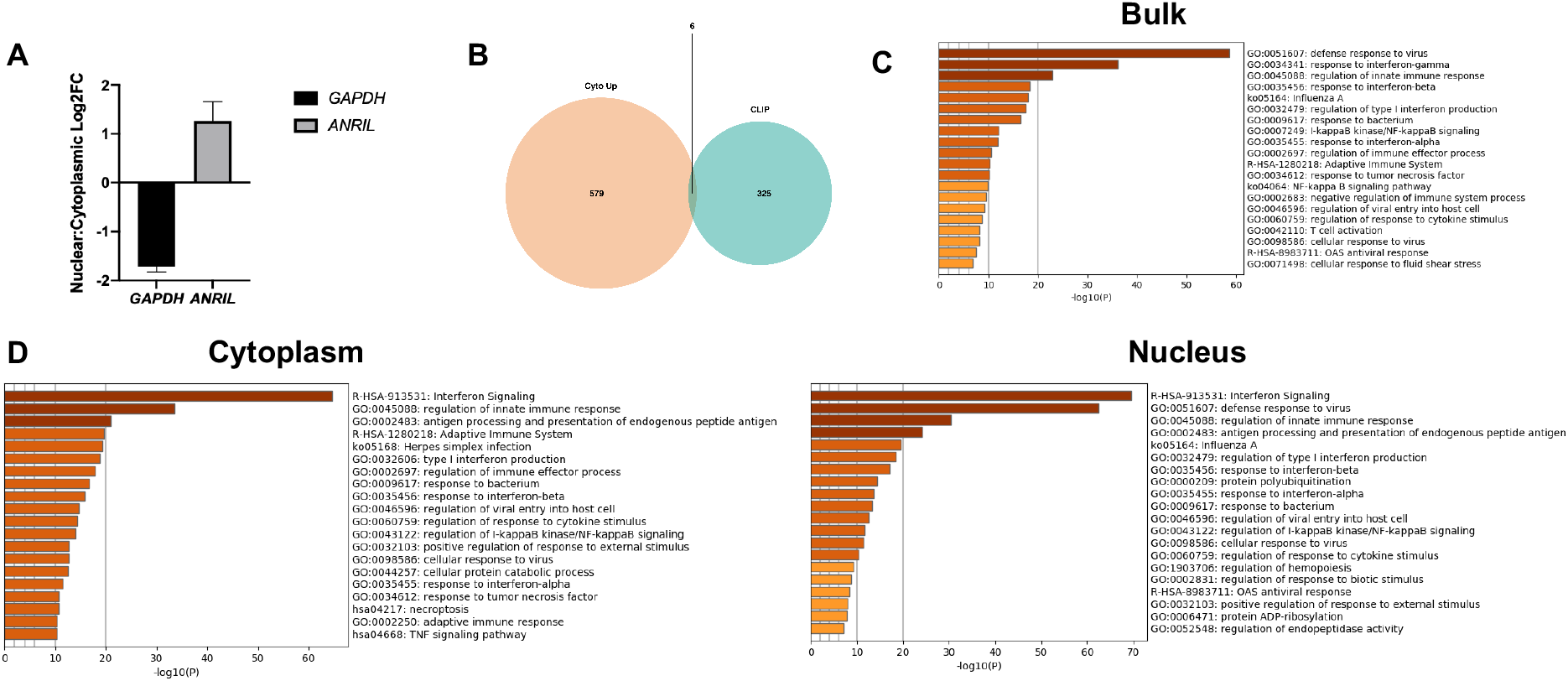
A) Log_2_ fold change from the fractionated RNA-sequencing between the nucleus and cytoplasm for markers of successful fractionation in the mock-infected samples: GAPDH for cytoplasm and ANRIL for nucleus. B) Comparison of transcripts increased in the cytoplasm and transcripts that lost interaction with UPF1 as determined by CLIP. Only 6 transcripts overlapped between the two groups. C) Metascape Analysis of significantly upregulated transcripts found in the bulk RNA sequencing of Figure 2A. The top 400 upregulated genes were used to produce this GO clustering. D) Metascape Analysis of significantly upregulated transcripts found in the cytoplasmic and nuclear fractionated RNA sequencing of Figure 2D. The top 400 and 300 upregulated genes were used to produce this GO clustering.

**Supplemental Figure 3.**
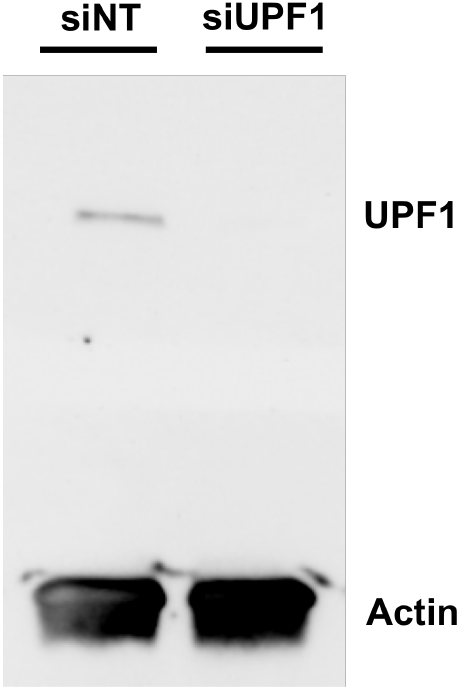
Western blot for UPF1 in siNT and siUPF1 treated cells. Actin is shown as a loading control.

**Supplemental Figure 4:**
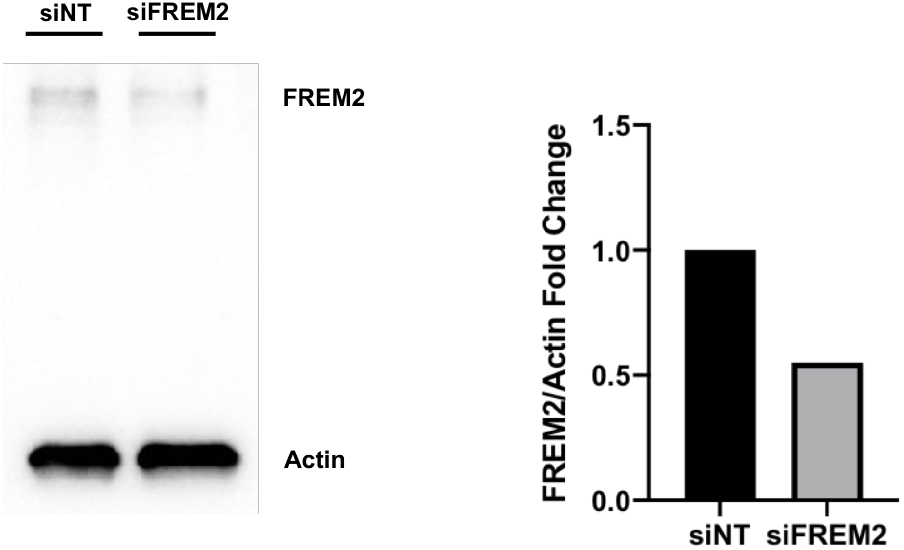
Western blot for FREM2 in siNT and siFREM2 treated NPCs after 7 days. Densitometric analyses of FREM2 were performed using ImageJ to quantify relative band intensities

## Notes

### Competing Interest Statement

The authors have declared no competing interest.

### Summary of Updates

Text was edited for clarity, author affiliations and funding sources updated.

